# Worldwide patterns of human epigenetic variation

**DOI:** 10.1101/021931

**Authors:** Oana Carja, Julia L. MacIsaac, Sarah M. Mah, Brenna M. Henn, Michael S. Kobor, Marcus W. Feldman, Hunter B. Fraser

**Affiliations:** Department of Biology, Stanford University, Stanford, CA; Centre for Molecular Medicine and Therapeutics, Child and Family Research Institute, University of British Columbia, Vancouver, BC, Canada; Department of Medical Genetics, University of British Columbia, Vancouver, BC, Canada; Department of Ecology and Evolution, Stony Brook University, Stony Brook, NY

**Keywords:** Illumina HumanMethylation450K Array, DNA methylation, RNA-seq, HGDP, PCA, population specificity, human diversity

## Abstract

DNA methylation is an epigenetic modification, influenced by both genetic and environmental variation, that can affect transcription and many organismal phenotypes. Although patterns of DNA methylation have been shown to differ between human populations, it remains to be determined whether epigenetic diversity mirrors the patterns observed for DNA polymorphisms or gene expression levels. We measured DNA methylation at 480,000 sites in 34 individuals from five diverse human populations in the Human Genome Diversity Panel, and analyzed these together with single nucleotide polymorphisms (SNPs) and gene expression data. We found greater population-specificity of DNA methylation than of mRNA levels, which may be driven by the greater genetic control of methylation. This study provides insights into gene expression and its epigenetic regulation across populations and offers a deeper understanding of worldwide patterns of epigenetic diversity in humans.

## Introduction

Human evolutionary history has left a strong signature on worldwide patterns of genetic variation (Cavalli-Sforza *et al.*, 1994; Li *et al.*, 2008; Novembre *et al.*, 2008). Principal component analyses and related methods have been instrumental in revealing how the genetic diversity of individuals varies within and across populations in exhibiting population stratification and admixture. The first two principal components of a single nucleotide polymorphism (SNP) genotype matrix are often sufficient to compare the ancestries of different populations and to show how genetic similarity between populations varies with geographic distance (Jakobsson *et al.*, 2008; Li *et al.*, 2008; Ramachandran *et al.*, 2005; Rosenberg *et al.*, 2005; Price *et al.*, 2006).

Here we ask whether this evolutionary history has left similar traces on worldwide patterns of epigenetic variation. The epigenome is situated at the interface between the genome and the environment, and their interactions may underlie the role of epigenetics in adaptation to the environment and other complex phenotypes. However, our understanding of the global epigenomic and transcriptomic diversity across human populations is far from complete (Heyn *et al.*, 2013; Fraser *et al.*, 2012; Martin *et al.*, 2014). Does epigenetic diversity exhibit signatures of human evolutionary history and do these signatures mirror the patterns of genetic variation? The relationship between the geographic patterns of ancestry in genomic and epigenomic variation is still uncharacterized in studies of genome-wide methylation in different human populations (Heyn *et al.*, 2013; Fraser *et al.*, 2012); however PCA on DNA methylation data from two populations did show partial separation (see Moen *et al.* (2013)). Previous studies aimed at understanding the patterns of diversity in gene expression levels have found that, unlike genotype-based PCA plots, PCs of expression data do not cluster by geographic location, and population ancestry cannot be determined using mRNA levels alone (see, for example, Stranger *et al.* (2012), Martin *et al.* (2014)).

Here, we analyze data on SNPs, CpG methylation levels and transcriptional variation (mRNA levels) for the same individuals from five different populations. These populations are among the Centre d’Etude du Polymorphisme Humain/Human Genome Diversity Panel (CEPH-HGDP) populations and span the breadth of human migration history (Cann *et al.*, 2002). Using this dataset, we compare the population specificity of the methylome and the transcriptome and study the correspondence between predefined population groups with those inferred from individual multi-locus genotypes and epigenotypes.

We find that DNA methylation and gene expression patterns of variation closely resemble the geographic population relationships inferred from the patterns of genomic variation. Small levels of differentiation between individuals at a large number of methylated sites are sufficient to cluster them into different groups that coincide with their ancestral origins. Moreover, we find that greater population specificity for the methylome than the transcriptome may be driven by tighter genetic control of CpG methylation. This finding provides further clues as to the contribution of the genome in shaping worldwide patterns of methylation and gene expression variation. Although our understanding of the establishment and maintenance of epigenomic variation is far from complete, these results provide a first resource for analyzing epigenetic population specificity and its genetic determination.

## Results

### Data

The data set comprises SNP, CpG methylation and mRNA sequencing information for 34 individuals from a sample of five of the CEPH-HGDP populations. The HGDP lymphoblastoid cell lines (LCLs) have been used extensively to study patterns of genetic variation (Cann *et al.*, 2002; Rosenberg *et al.*, 2002; Li *et al.*, 2008). The five populations were chosen to capture differences in genetic diversity that stem from serial founder effects throughout human evolutionary history (Ramachandran *et al.*, 2005). The 34 individuals in the study include six Yakut, seven Cambodian, seven Pathan, seven Mozabite and seven Maya individuals. Geographic locations of the samples were previously reported by Cann *et al.* (2002).

The genotype data used were previously reported (Li *et al.*, 2008) and 644, 258 markers passed our quality control filters and were kept for subsequent analyses. Genome-wide methylation patterns were assessed using the Illumina 450K Methylation array (Sandoval *et al.*, 2011) which quantifies methylation at 480,000 CpG sites genome-wide. After extensive normalization and quality control (see **SI: Methods and Materials**), the data used in the analyses here consist of 317, 109 CpG sites. The mRNA data used consists of expression abundance levels determined using cufflinks-2.0.2; the FPKM (fragments per kilobase of exon per million mapped reads) estimates per transcript were previously reported in Martin *et al.* (2014).

### Context of population genetic variation

Worldwide patterns of allele frequencies reflect geographic variation in demographic structure and adaptation to environmental differences among different populations. Genetic variation has been shown to closely correspond to self-identified groups or to geographically and linguistically similar populations (Cavalli-Sforza *et al.*, 1988; Ramachandran *et al.*, 2005; Conrad *et al.*, 2006; Jakobsson *et al.*, 2008; Creanza *et al.*, 2015). This general agreement between genetic variation and geographic location has also been extensively studied using the HGDP dataset (Rosenberg *et al.*, 2002; Li *et al.*, 2008).

The genetic context for the five populations used in this paper is provided in **Figure 1**. **Figure 1A** shows the geographic locations of our samples, as previously reported in Cann *et al.* (2002). We performed principal component analysis (PCA) on the identity-by-state (IBS) genotype matrix to visualize the patterns of genetic variability. The first and second PCs explain 26% and 24% of the IBS variation, respectively, and clearly differentiate the individuals into five well-separated clusters that correspond to the five populations sampled (**Figure 1B**). Even with the limited sample size, the population structure revealed by the SNP frequencies is extremely robust. To facilitate comparison between the genetic and epigenetic datasets we quantify the strength of the genomic PCA clustering by computing the Silhouette scores (Lovmar *et al.* (2005) and **SI: Methods and Materials**) for the SNPs in the five populations as well as the average Silhouette for the entire data set (**Supplementary Figure S1**). The Silhouette score of an individual measures how similar it is to its own predefined population cluster, relative to individuals in other clusters. The average Silhouette cluster score across all individuals is a measure of how tightly the data can be assembled into population clusters. The average Silhouette score for the genetic clustering presented in **Figure 1B** is 0.823. A tree generated using hierarchical clustering also captures the genetic relationships between the individuals and their populations (**Figure 1C**). The branching pattern of this tree agrees with the accepted order of ancestral human expansion, consistent with the out of Africa hypothesis (Cavalli-Sforza and Feldman, 2003; Ramachandran *et al.*, 2005; Henn *et al.*, 2012).

**Figure 1.**
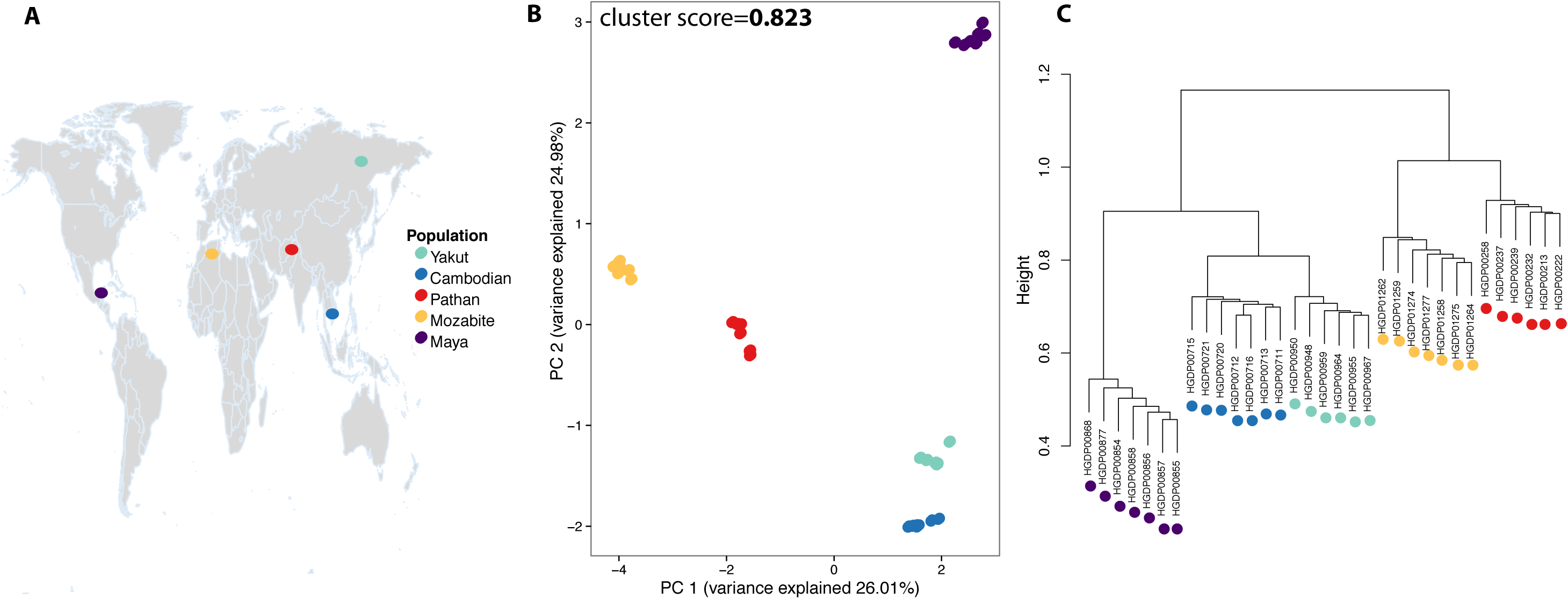
Context of genome-wide population structure. **Panel A**: the geographic locations of populations in the dataset. **Panel B**: PCA on SNP IBS matrix. The first and second PCs explain 26% and 24% of the IBS variation, respectively, and clearly differentiate the individuals into five well-separated clusters that correspond to the five populations sampled. **Panel C**: A Hierarchical clustering tree also captures the genetic relationships between the individuals and their populations.

### Population specificity of CpG methylation

We quantified patterns of population specificity at the methylation level and computed the number of CpG sites with DNA methylation differences between the five different populations studied.

For every CpG site, we used the Kruskall-Wallis (K-W) test to quantify population difference, assign *p*-values and identify CpG sites that are significantly differentially methylated between the five different populations. **Figure 2A** shows that there exist significant differences in CpG methylation between the five populations compared to the uniform *p*-value distribution that is expected by chance (black line). We find 7084 CpG sites with a K-W *p*-value smaller than 0.01 (24% FDR), 321 CpG sites with a K-W *p*-value smaller than 0.001 (12% FDR) and four CpG sites with a K-W *p*-value smaller than 0.0001 (3% FDR).

**Figure 2.**
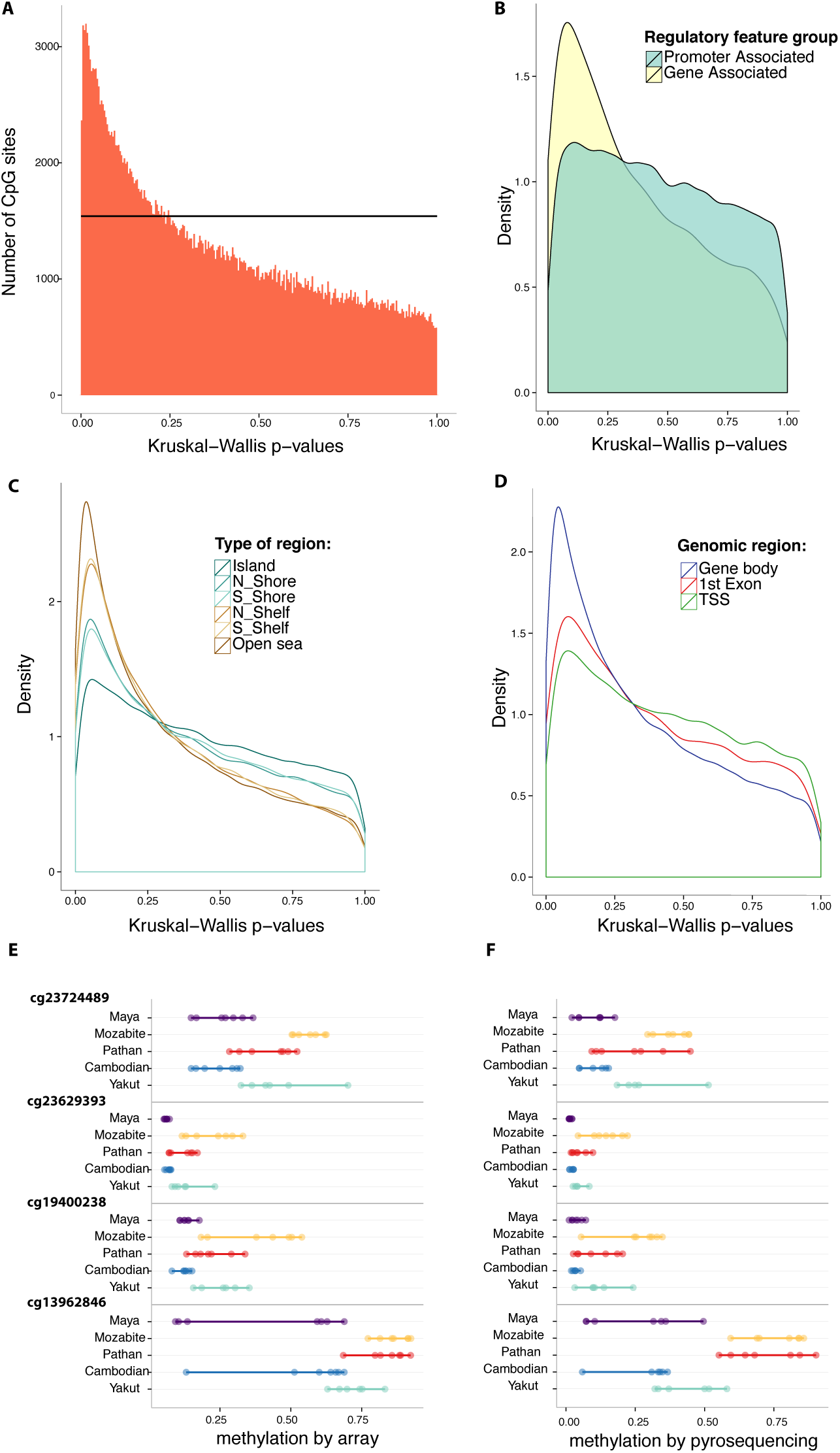
Population specificity of CpG methylation. **Panel A**: Graph of KruskalWallis *p*-values for all CpG markers across all individuals in the five different populations. The black horizontal line corresponds to the uniform *p*-value distribution expected by chance. Differences based on different types of CpG regions are plotted in **Panels B**, **C** and **D**. The CpG sites that exhibit population differentiation are enriched in regions that are gene-associated, outside of genomic islands, and inside gene bodies. **Panels E** and **F**: Comparison of percentage methylation by array (**E**) and by pyrosequencing (**F**) for the top four CpG sites with highest population specificity.

Using information on the genomic location of the CpG sites, we investigated how these population differences might vary across different genomic regions (**Figure 2B-D**), correcting for differences in the number of interrogated CpG sites across regions. We found that population differences are enriched in CpG islands, which are promoter regions with high CpG content.

These population-specific sites decrease in frequency in regions flanking the islands, the CpG Shores and the CpG Shelves. **Figure 2** also shows that population-specific sites are enriched in gene bodies.

We replicated the results for the four CpG sites with K-W *p*-value smaller than 0.0001 by pyrosequencing bilsufite-treated DNA. The results show excellent concordance between the K-W p-values obtained using the Illumina array and those obtained by pyrosequencying (**SI, Table S1**). The percentage methylation by array and by pyrosequencing is shown in **Figures 2E-F**. The average differences in methylation as well as direction of level of CpG methylation across the five populations are also conserved between the two measurements.

### Structure of epigenetic population variation

We measured epigenetic divergence by computing *P*_*st*_, the phenotypic differentiation between populations (Pujol *et al.*, 2008; Edelaar *et al.*, 2011; Leinonen *et al.*, 2013), for methylation and expression sites across the genome. This measure is analogous to *F*_*st*_ (Weir and Cockerham, 1984), varies from 0 to 1, and estimates population differentiation for quantitative traits. For a given epigenetic mark, 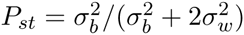, where 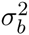 is the between population variance and 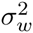 is the average within population variance (see **SI:Methods and Materials**).

Selecting the top 0.5% of CpG sites that exhibit the highest population divergence in methylation (i.e., the highest *P*_*st*_ values), we performed principal component analyses to assess patterns of epigenetic variability between the five different populations. We repeated this analysis using the same number of mRNA expression markers that exhibit the highest population divergence. Using only the most differentiated markers, the methylation levels cluster individuals by population (**Figure 3** left panel), and the clustering patterns are similar to those obtained from SNP data. The mRNA expression data, presented in **Figure 3**, right panel, exhibits similar clustering patterns. The CpG methylation Silhouette cluster score is higher than that for expression, as also observed from the average Silhouette scores (Silhouette plots are presented in **Supplementary Figures S4** and **S5**. The results of an equivalent analysis using the markers with the smallest K-W *p*-values are presented in **Supplementary Figure S2**. The Silhouette cluster score as a function of number of the top markers used is presented in **Supplementary Figure S6**. Using only a small number of population-specific epigenetic marks (CpG methylation and mRNA expression marks) we recapture signatures of human evolutionary history on world-wide population structures that were previously observed at the genetic level. The epigenetic similarity varies with geographic distance and is surprisingly consistent with previously hypothesized human migration patterns out of Africa.

**Figure 3.**
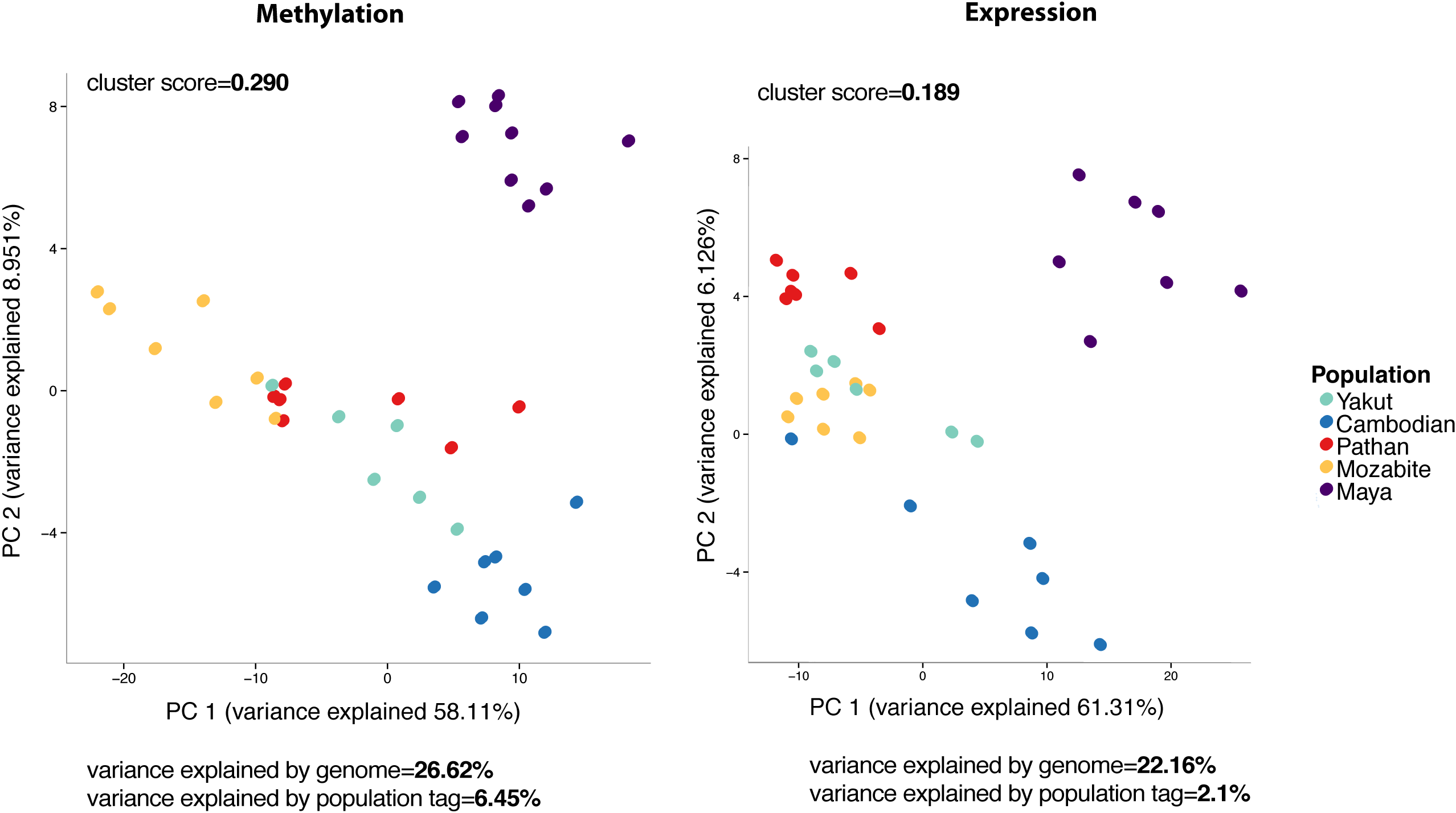
Structure of epigenome-wide population differences. Left: PCA using top 200 CpG sites with highest *P*_*st*_ values. Right: PCA using top 200 mRNA expression sites with highest *P*_*st*_ values. Silhouette cluster scores and percentage of variance explained by the genome versus the population tag are as presented.

### Drivers of the observed epigenetic population variation

The observed epigenetic differences between populations could be caused by genetic or environmental variation, or a combination of both. Since these data are from lymphoblastoid cell lines (LCLs) that were grown in a controlled laboratory environment, the more likely driver of the observed differences is the genetic background. For example, both CpG methylation and mRNA levels could be influenced by between-population differences in allele frequencies at genetic variants that control the epigenome. To investigate how much of the observed population specificity can be explained by genetic variation, we first identified the local SNP (in a 200kb window from the CpG site, or the transcription start site (TSS) for mRNA) most strongly associated with each of the top 0.5% most variable CpGs or mRNA levels across all of our samples. We then performed an analysis of variance including these single SNP genotypes for each of the epigenetic marks used in **Figure 3** to assess whether the SNPs genotype or population was a stronger predictor of methylation and expression (**SI: Materials and Methods**). We computed the average variance explained by genotype versus the population label across all the markers used for the PCA in **Figure 3**. We found that the sites with the highest degrees of population specificity were more strongly associated with the local SNP than with population, and this local SNP explained a much higher percentage of the variance than the population label (the SNP genotype explained 26% of the variance for methylation and 22% for expression, while the population label explained 6% for methylation and 2% for expression). The higher Silhouette cluster score using CpG methylation levels compared to mRNA expression levels might therefore be explained by a larger degree of genetic control (**Figure 3**). This result also indicates that cell line artifacts are most likely not responsible for the population specificity we observe since they would be unlikely to correlate with the SNP genotypes (Fraser *et al.*, 2012).

## Discussion

Characterization of the variability of the human epigenome is essential for investigating the mapping from genotype to phenotype as well as the role of the epigenome in diseases. The establishment and maintenance of CpG methylation is controlled by many factors and the relative importance of stable genetic control (Bell *et al.*, 2011; Fraser *et al.*, 2012; Gibbs *et al.*, 2010; Schalkwyk *et al.*, 2010) versus plastic environmental influence (Breitling *et al.*, 2011; Jirtle and Skinner, 2007; Feil and Fraga, 2012) remains unclear. Similarly, transcriptional diversity reveals both stable gene expression levels regulated by genetic variation (Zhang *et al.*, 2008), as well as associations with numerous environmental exposures (Jaenisch and Bird, 2003).

Using high resolution maps of genome-wide CpG methylation and RNA sequencing, we have analyzed worldwide patterns of methylation and expression level variation across five populations selected for their geographic diversity (Cavalli-Sforza and Feldman, 2003; Ramachandran *et al.*, 2005). Our data allow a characterization of human epigenetic variation and its comparison to human genetic variation. While cell culture can induce epigenetic changes in LCLs, existing variation between different individuals is typically preserved (Caliskan *et al.*, 2011).

We used Principal Component Analysis (PCA) to explore the relative patterns of methylation and transcription diversity and compare them with genetic patterns of variation. Despite their limitations for inferring the underlying causal processes that give rise to the PCA plot (Novembre and Stephens, 2008), they remain a useful tool for exploring and describing population substructure in analyses of population genetic variation. The Silhouette scores for the clustering of individuals in PC space is much higher for genetic data than those for epigenetic data, however the patterns of variation across the five populations are preserved.

Epigenetic divergence across populations reveals higher population specificity of CpG methylation than mRNA expression. Our results demonstrate stronger genetic control of interpopulation CpG methylation levels than of corresponding mRNA expression levels, with likely downstream consequences. Because of the small sample size, we cannot provide an accurate estimate of how much of the population-specific DNA methylation we observed is due to global mSNPS (SNPs that control methylation levels (Bell *et al.*, 2011; Fraser *et al.*, 2012)), population specific mSNPs, or differences in environment, including genotype-by-environment interactions.

Through the accumulation of small allele-frequency differences across many loci, previous studies have identified geographic patterns from allele frequency variation among human populations (Rosenberg *et al.*, 2005; Ramachandran *et al.*, 2005). Understanding patterns of worldwide human epigenetic diversity is also essential for understanding how population substructure can shape the architecture of phenotypic traits. This epi-structure of human populations could be particularly relevant in various medical contexts. Variation in both genetic and epigenetic disease phenotypes, risk factors for different environmental exposures or differences in drug response may depend on ancestry and be population specific (Jirtle and Skinner, 2007; Feinberg, 2007). Further thorough characterizations of worldwide human epigenetic variation will likely prove to be informative for understanding the origins of human phenotypic variation.

## Acknowledgements

OC and MWF acknowledge support from the Morrison Institute for Population and Resource Studies at Stanford and the Stanford Centre for Computational, Evolutionary and Human Genomics. The authors and BMH acknowledge support from NIH grant 3R01HG003229 to Carlos B. Bustamante. MSK is a Senior Fellow of the Canadian Institute for Advanced Research and the Canada Research Chair in Social Epigenetics. We thank members of the Feldman Laboratory, in particular Nicole Creanza and Julie M. Granka, for helpful discussions and Meaghan Jones for comments on an initial draft.

---

**Figure S1.**
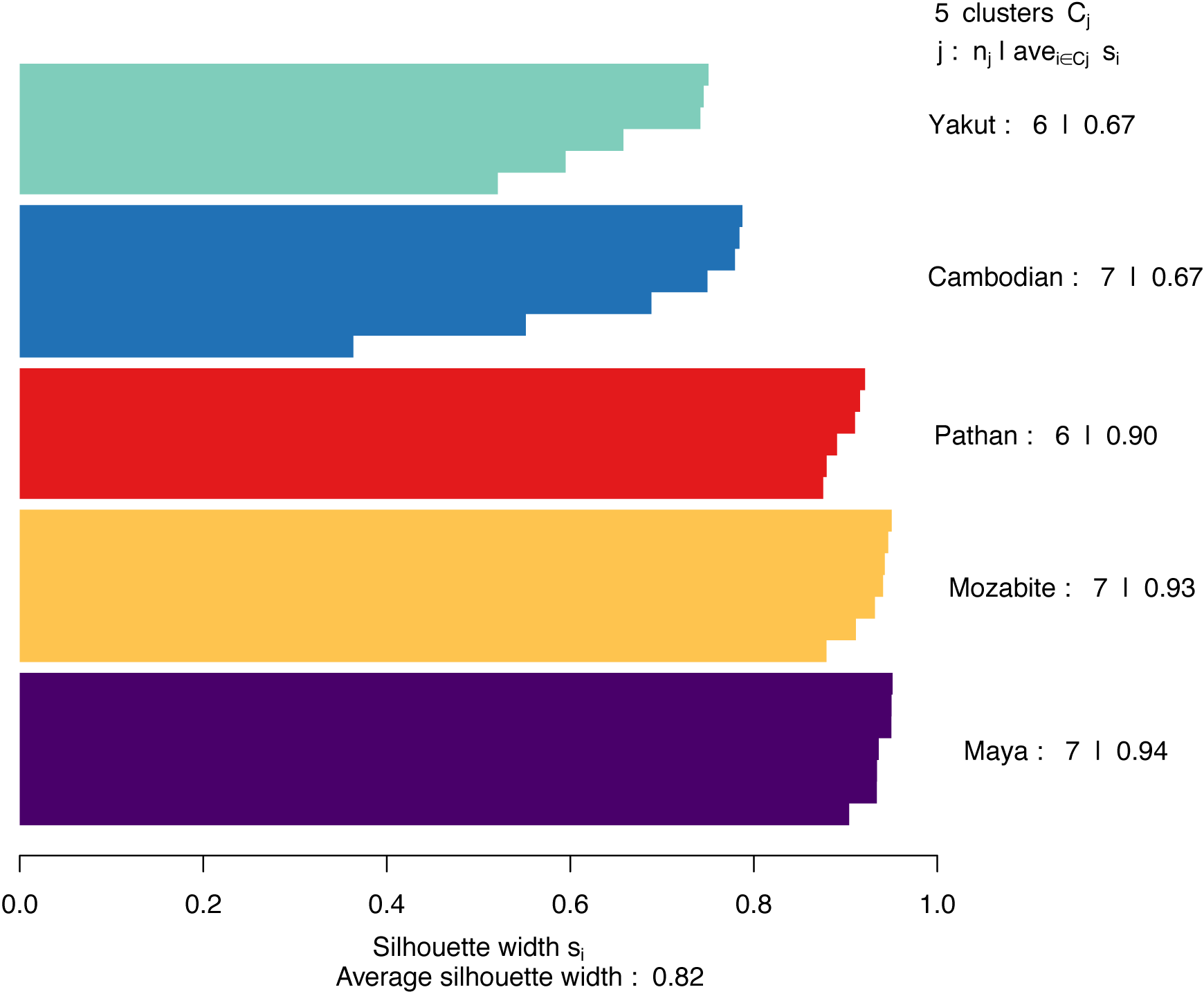
Silhouette plots using SNPs. Average Silhouette score is 0.82.

**Figure S2.**
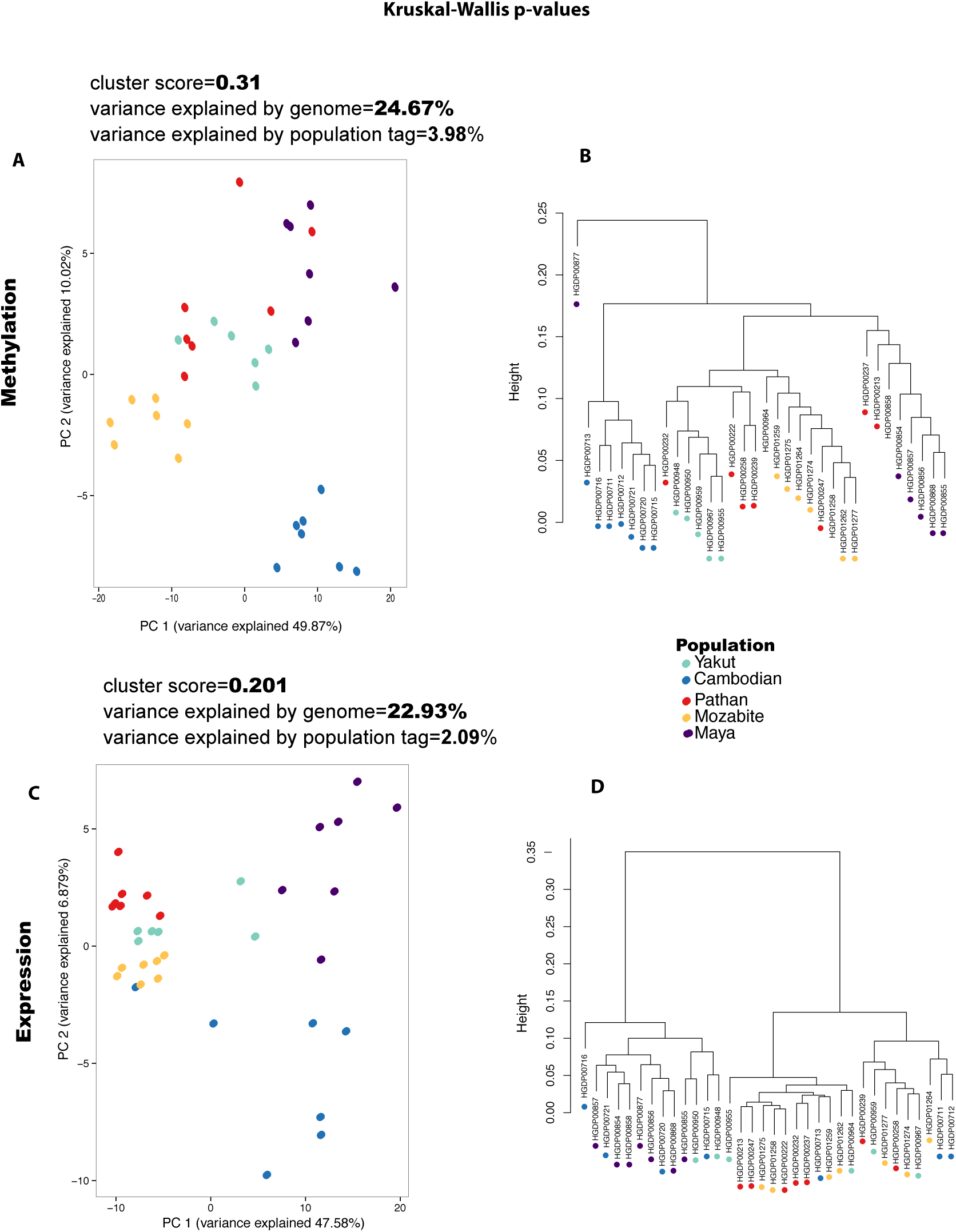
PCA plots and hierarchical clustering trees using top 200 smallest KruskalWallis *p*-values for CpG methylation and mRNA expression markers. **Panel A**: PCA plot using CpG methylation levels. **Panel B**: Hierarchical clustering tree using CpG methylation levels. **Panel C**: PCA plot using mRNA expression levels. **Panel D**: Hierarchical clustering tree using mRNA expression levels.

**Figure S3.**
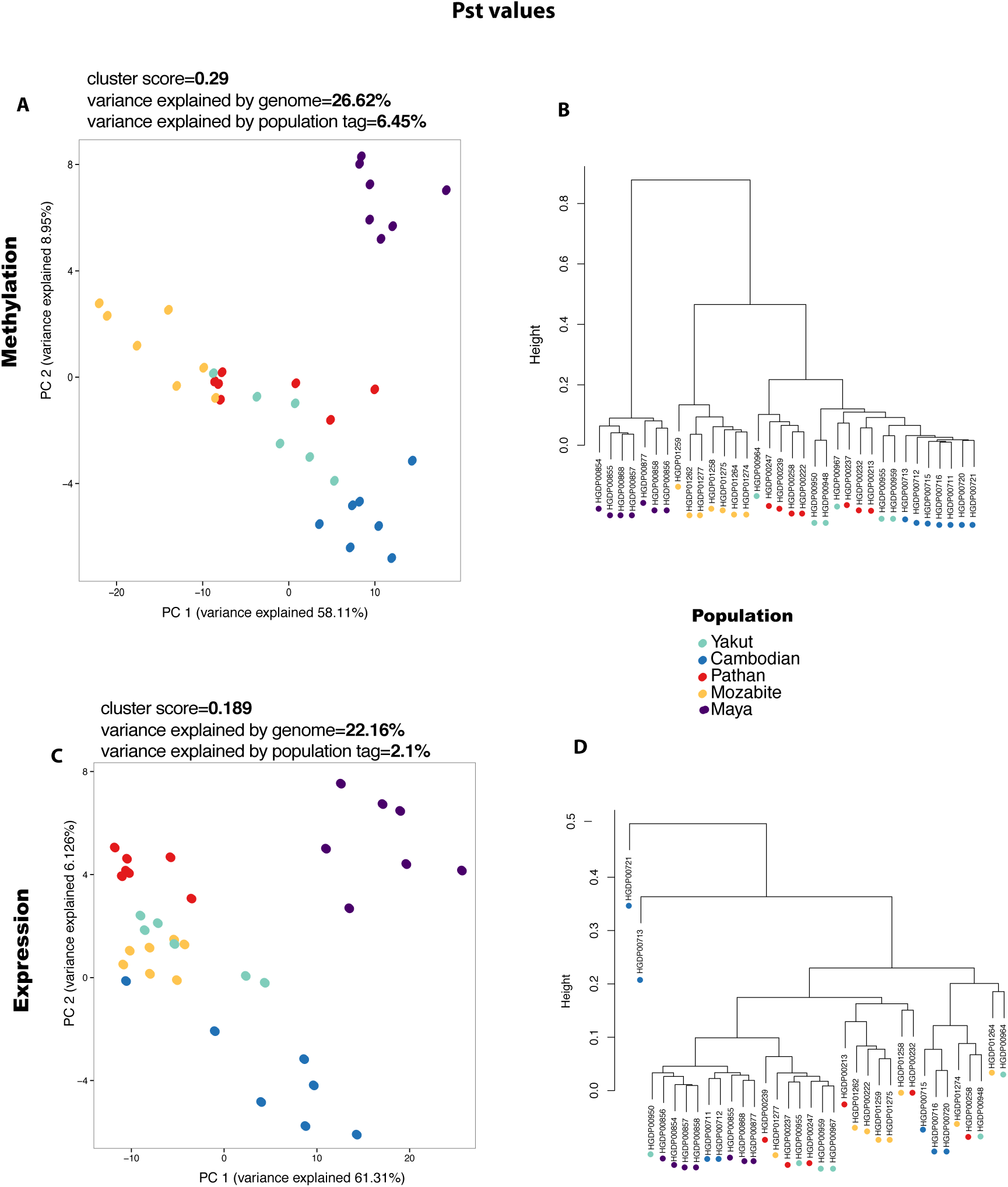
PCA plots and hierarchical clustering trees using top 200 largest *P*_*st*_ values for CpG methylation and mRNA expression markers. **Panel A**: PCA plot using CpG methylation levels. **Panel B**: Hierarchical clustering tree using CpG methylation levels. **Panel C**: PCA plot using mRNA expression levels. **Panel D**: Hierarchical clustering tree using mRNA expression levels.

**Figure S4.**
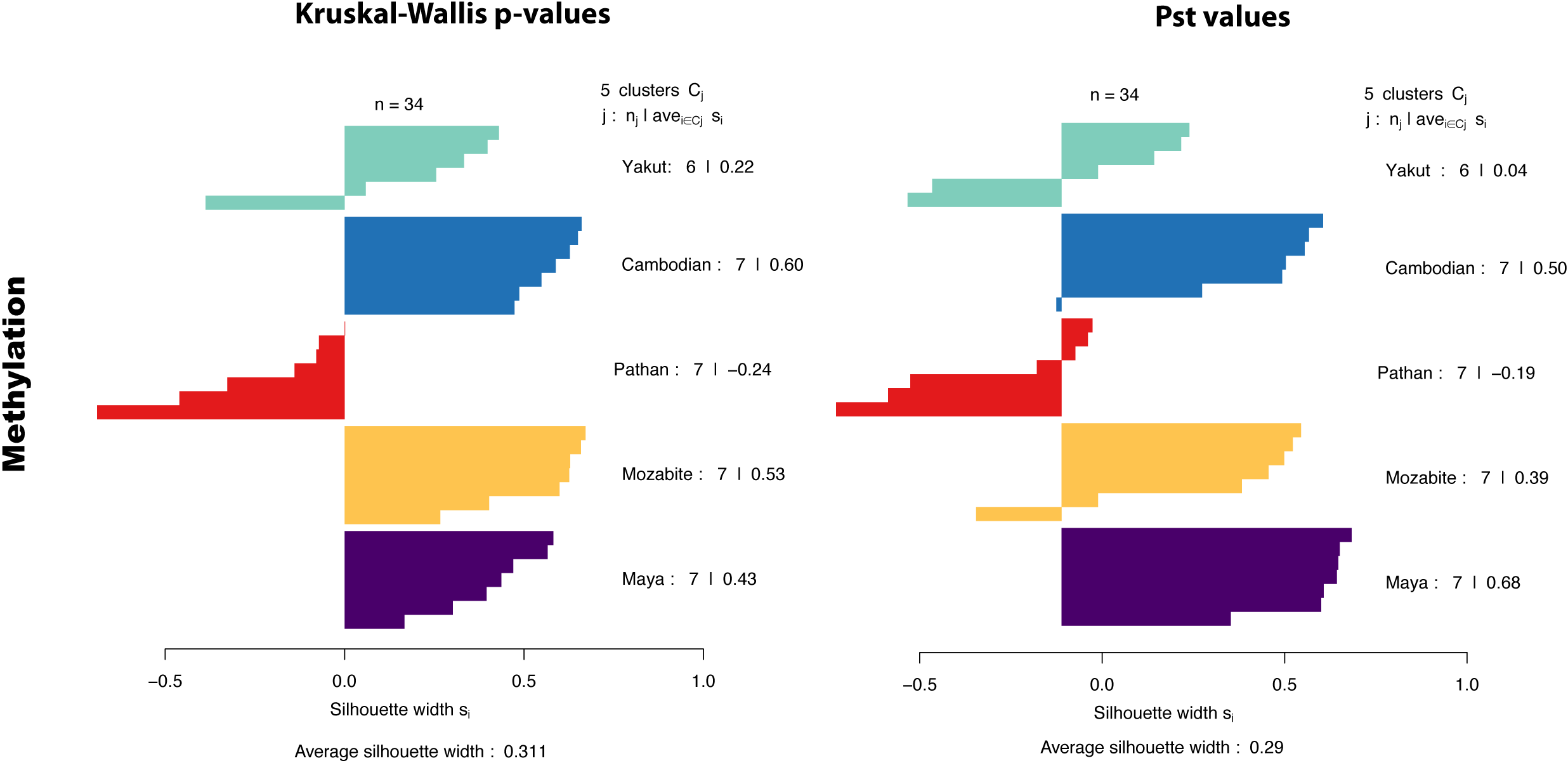
Silhouette plots using the top methylation markers.

**Figure S5.**
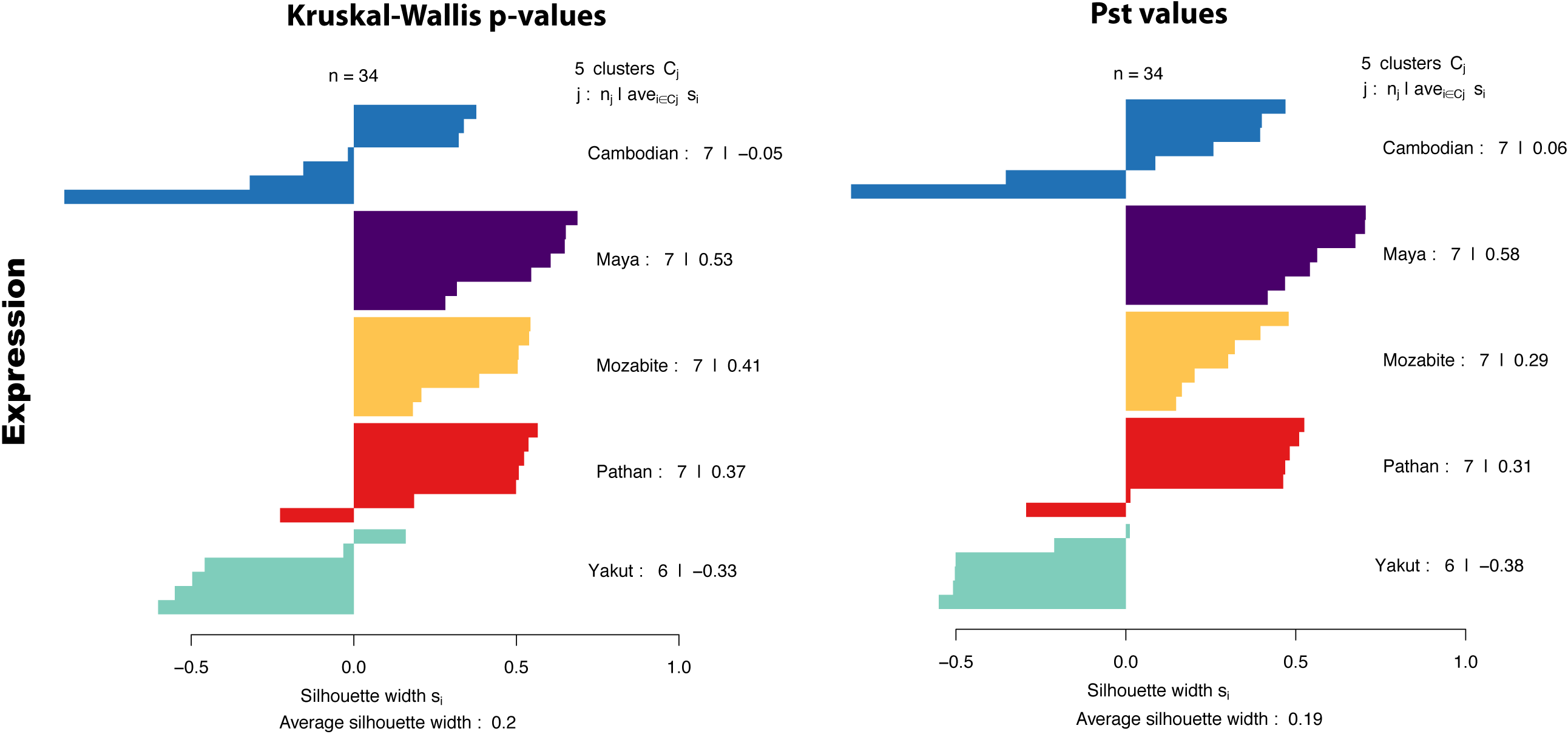
Silhouette plots using the top mRNA expression markers.

**Figure S6.**
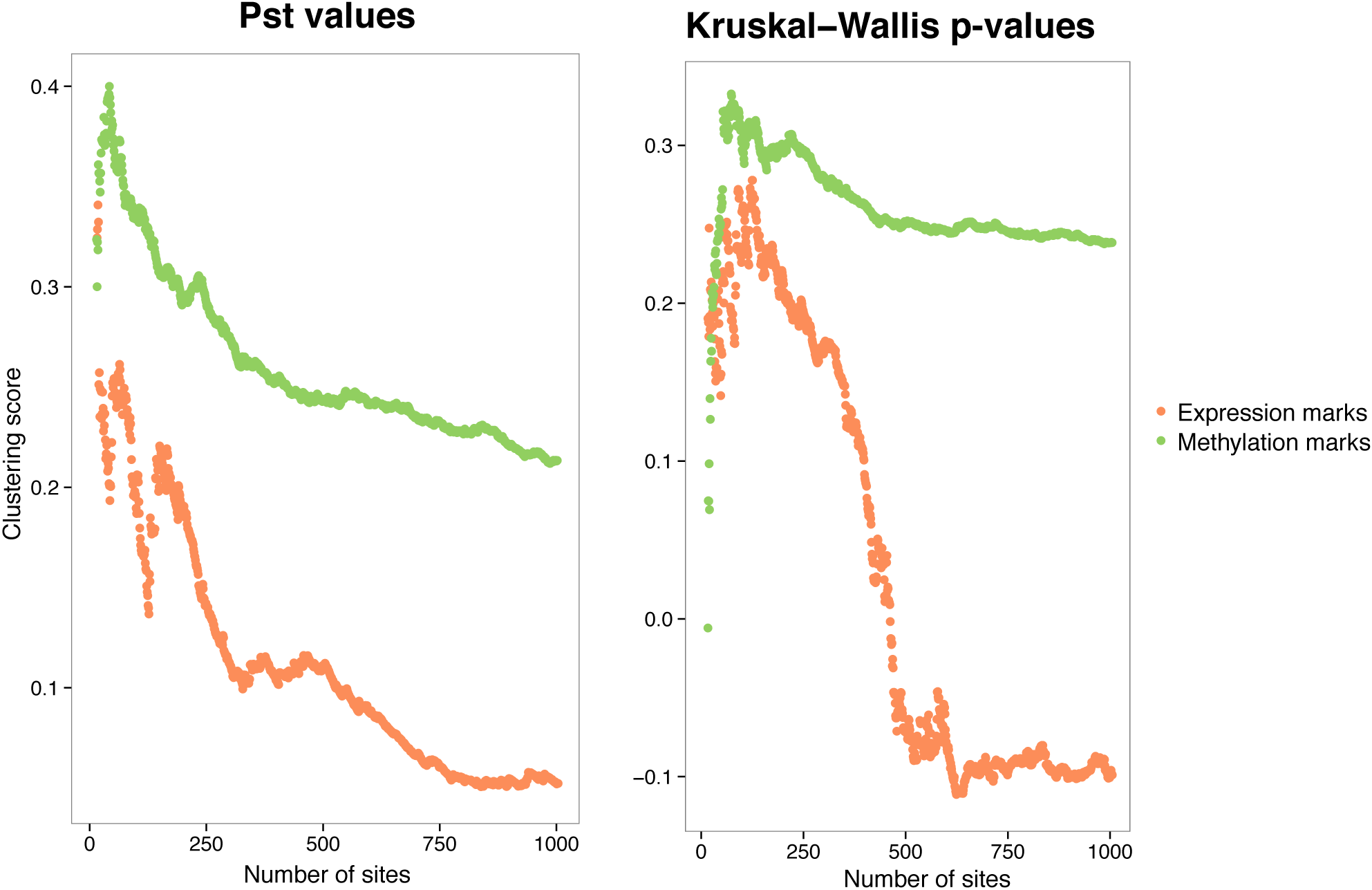
Silhouette cluster score as a function of number of markers used.

**Figure S7.**
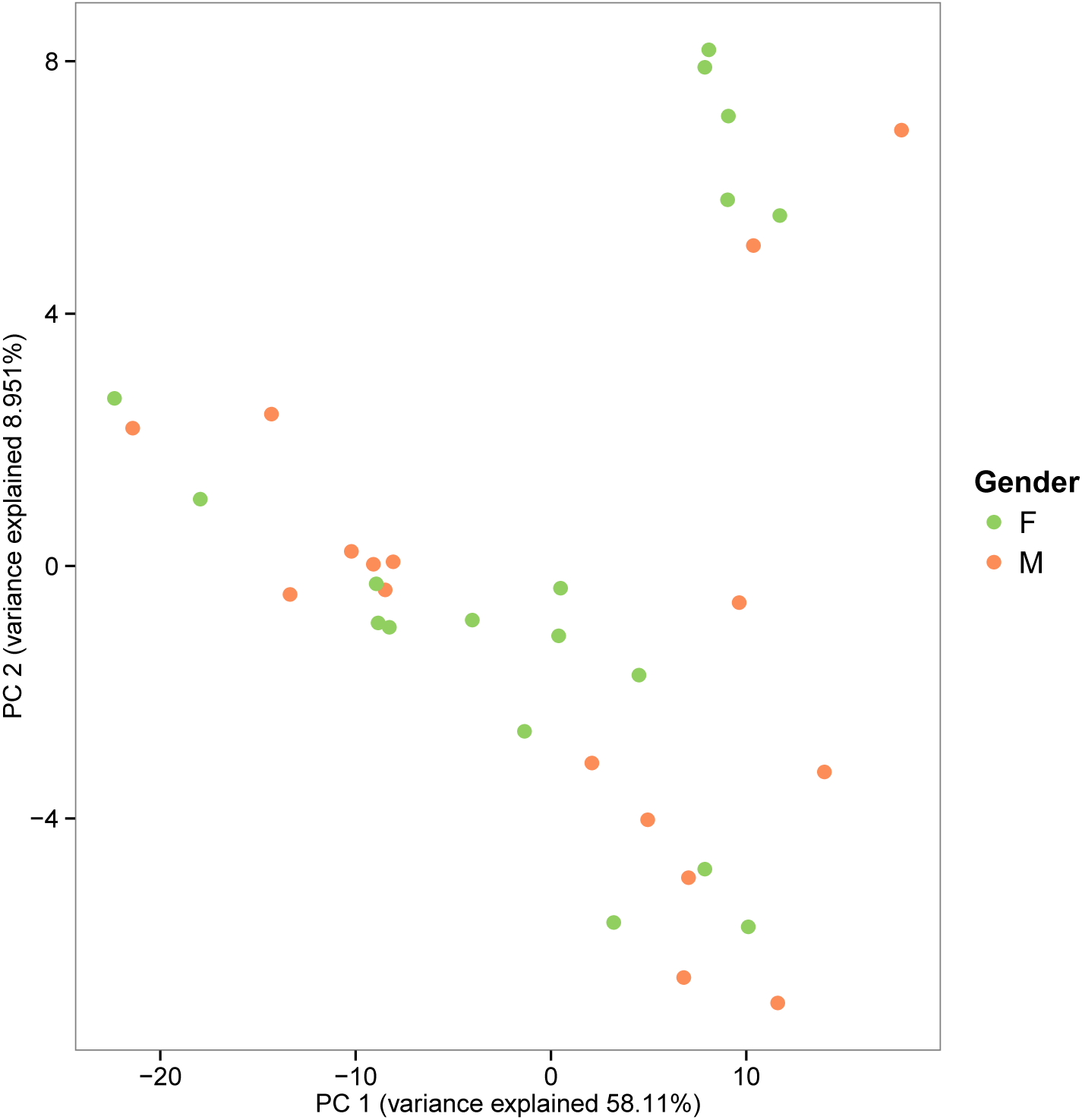
Clustering not due to a gender effect. PCA using top 200 CpG sites with highest *P*_*st*_ values (same as **Figure 3**). Individuals are no longer labeled by population, but by gender. The clustering patterns observed are therefore not driven by a gender effect.

## SI: Materials and Methods

### Samples

Individuals were selected from the HGDP-CEPH Human Genome Diversity Cell Line Panel: 6 Yakut individuals, 7 Cambodian individuals, 7 Pathan individuals, 7 Mozabite individuals, and 7 Maya individuals. The geographic locations of these populations were previously reported in Cann *et al.* (2002).

### Silhouette scores

The Silhouette value for each point is a measure of how similar that point is to points in its own cluster, when compared to points in other clusters. In our case, the clusters are the populations. The Silhouette value for the *i*-th point, *S*_*i*_, is defined as *S*_*i*_ = (*b*_*i*_ - *a*_*i*_)*/max{a_i_, b_i_}*, where *a*_*i*_ is the average distance from the *i*-th point to the other points in the same cluster as *i*, and *b*_*i*_ is the minimum average distance from the *i*-th point to points in a different cluster, minimized over clusters. The silhouette value ranges from -1 to 1 and a high Silhouette value indicates that *i* is well-matched to its own population, and poorly-matched to neighboring populations. If most individuals have a high Silhouette value, then the clustering solution is appropriate. If many individuals have a low or negative Silhouette value, then the clustering solution may have either too many or too few populations. The Silhouette clustering evaluation criterion was used with the Euclidean distance, but can be used with any distance metric.

### Genome-wide human DNA methylation data

DNA methylation measurements of bisulfite-treated genomic DNA were performed with the HumanMethylation450 BeadChip assay (Illumina, San Diego, CA, USA), quantifying methylation at 485,000 sites per sample at singlenucleotide resolution, using experimental procedures recommended by the manufacturer. The bisulfite-converted DNA is subjected to a whole-genome amplification step, followed by fragmentation and hybridization to probes on the microarray. Following hybridization, allele-specific single-base extension of the probes incorporates a fluorescent label (ddNTP) for detection. Using the Illumina GenomeStudio software provided by the manufacturer, methylation levels (*β* values) were then computed by dividing the methylated probe signal intensity by the sum of methylated and unmethylated probe signal intensities. These *β* values range from 0 (completely unmethylated) to 1 (completely methylated) and provide a quantitative readout of relative DNA methylation for each CpG site within the whole cell population. Samples from the five populations were run together in a randomized order to avoid confounding batch effects with population differences. Technical replicates across different runs had correlation *r >* 0.99. All our samples passed internal controls included on the HumanMethylation450 array, including controls for array background, hybridization quality, target specificity and bisulfite conversion. Furthermore, all samples passed quality control check of having detection P-value *>* 0.05. Subsequent cluster analysis indicated the absence of any outlier samples.

### Normalization of *β* values across individuals

The data were color corrected, background corrected, quantile normalized and SWAN normalized to correct for type I and type II difference (Maksimovic *et al.*, 2012). To perform the background normalization, background intensity, as measured by negative background probes present on the array, was subtracted from the raw intensities to adjust for varying background signals across different samples. This background adjustment was done separately for raw data from the green and red channels to adjust for Cy3 and Cy5 differences. All negative intensities were assigned values of zero before further normalizations were performed. To minimize batch effects across different sets of arrays, background adjusted raw data from both channels were quantile-normalized separately. The quantile normalization is done at the intensity level, whereas the SWAN normalization is done at the *m*-value level and includes a step which randomly chooses a subset of type II probes to normalize to type I probes and then normalizes the rest of the type II probes to the normalized type II probes. This randomization step results in slightly different result every time SWAN normalization is done, so in comparing *β* values created from one normalization run to those in another, it is usual to see slight differences. The*β* values were obtained after obtaining the *m*-values, using the formula *β* = 2^*m*^/(1 + 2^*m*^). In all of our analyses, we used *β* values since we saw no differences in the genome-wide trends or the top sites when using *m*-values. We prefer *β* values because they seem easier to interpret.

After quality control check, normalization and filtering out markers overlapping known SNPs in the 1, 000 Genome database and markers on the sex chromosomes, the CpG methylation data consists of *β* values for 317, 109 CpG sites.

### Calculation of false discovery rates (FDRs)

The FDRs were computed by permutation, which preserves aspects of the data that might affect the results of the analyses. For the population-specific methylation analysis, the FDRs were estimated using 1000 randomizations where the population tags were assigned randomly to every individual and the Kruskal-Wallis *p*-values were recomputed on this randomized data.

### Validation of population specific CpG sites through bisulfite pyrosequencing

Bisulfite PCR-pyrosequencing assays were designed with PyroMark Assay Design 2.0 (Qiagen). The regions of interest were amplified by PCR using the HotstarTaq DNA polymerase kit (Qiagen) as follows: 15 minutes at 95C (to activate the Taq polymerase), 45 cycles of 95C for 30s, 58C for 30s, and 72C for 30s, and a 5 minute 72C extension step. For pyrosequencing, a single-stranded DNA was prepared from the PCR product with the Pyromark Vacuum Prep Workstation (Qiagen) and the sequencing was performed using sequencing primers on a Pyromark Q96 MD pyrosequencer (Qiagen). The quantitative levels of methylation for each CpG dinucleotide were calculated with Pyro Q-CpG software (Qiagen). Primer sequences are available upon request.

### Concordance between array and pyrosequencing percentage methylation

For the four most differentiated sites, the K-W p-values using the Illumina array and the ones obtained by pyrosequencying are presented in table **S1** below.

**Table S1.** Comparing K-W p-values between array and pyrosequencing.

| Site | Array K-W p-value | Pyrosequencing K-W p-value |
| --- | --- | --- |
| cg13962846 | 5.065e-05 | 5.216e-05 |
| cg19400238 | 7.736e-05 | 0.0005818 |
| cg23629393 | 8.958e-05 | 9.736e-05 |
| cg23724489 | 8.688e-05 | 0.0005995 |

### The *P*_*st*_ values

*P*_*st*_ is a measure of the proportion of variance explained by between-population divergence. It is the phenotypic analog of the population genetics parameter *F*_*st*_ (Leinonen *et al.*, 2013; Pujol *et al.*, 2008). For a single probe, *P*_*st*_ was calculated as: 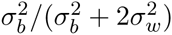, where 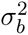 is the between population variance and 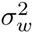 is the average within population variance. *P*_*st*_ values range from 0 to 1, with values near 1 signifying that the majority of epigenetic variance for a probe is between populations rather than within populations.

### Analysis of variance using local SNPs

For every CpG and mRNA marker, the local SNP was defined as the SNP within a 200kb window from the CpG site or the transcription start site (TSS) of the mRNA marker with the largest correlation with the marker levels across all individuals. We restricted our analysis to the 200 CpG and mRNA markers used in **Figure 3**. We performed an analysis of variance to obtain the variance explained by the population tag and the genetic marker for every population-specific epigenetic marker: *marker*_*level*_ *∼ P opulation* + *SN P*, where *marker*_*level*_ denotes the methylation opt expression level of that marker, *P opulation* denotes the population tag of the individual and *SN P* denotes the genotype of the individual. We then averaged the variances across the markers to obtain the average variance presented in **Figure 3**.

